# Cardiac valve regeneration in adult zebrafish: importance of TGFß signaling in new tissue formation

**DOI:** 10.1101/697706

**Authors:** Anabela Bensimon-Brito, Srinath Ramkumar, Giulia L. M. Boezio, Stefan Guenther, Carsten Kuenne, Héctor Sánchez-Iranzo, Dijana Iloska, Janett Piesker, Soni Pullamsetti, Nadia Mercader, Dimitris Beis, Didier Y. R. Stainier

**Author notes:** These authors contributed equally (Srinath Ramkumar, Giulia L. M. Boezio).

## Abstract

Cardiac valve disease can lead to severe cardiac dysfunction and is thus a frequent cause of morbidity and mortality. Its main treatment is valve replacement, which is currently greatly limited by the poor recellularization and tissue formation potential of the implanted valves. As we still lack suitable animal models to identify modulators of these processes, here we used the adult zebrafish and found that, upon valve decellularization, they initiate a striking regenerative program that leads to the formation of new functional valves. After injury, endothelial and kidney marrow-derived cells undergo cell cycle re-entry and differentiate into new extracellular matrix-secreting valve cells. The Transforming Growth Factor beta (TGFβ) signaling pathway promotes this process by enhancing progenitor cell proliferation as well as valve cell differentiation. These findings reveal a key role for TGFβ signaling in valve regeneration and also establish the zebrafish as a model to identify and test factors promoting valve recellularization and growth.

## Introduction

Cardiac valve disease is a major threat to human health worldwide, ultimately leading to heart failure and death. Valve defects affect 1% of all newborns but can also appear later in life with an incidence of 13.3% in people older than 75 years (Iung, 2003; Nkomo, et al., 2006; Yutzey, et al., 2014). The majority of diseased valves are not repairable, and the only possible therapy relies on surgical replacement to improve survival and quality of life (van Geldorp, et al., 2013). Due to the obvious lack of biological material, the field has invested in finding alternatives for implants, such as the use of mechanical, synthetic and biological scaffolds (Neuenschwander and Hoerstrup, 2004; Fallahiarezoudar, et al., 2015; Ibrahim, et al., 2017; Goecke, et al., 2018; Motta, et al., 2019). The growth and stability of these scaffolds upon implantation rely on the efficient recellularization of the matrix (VeDepo, et al., 2017; Bouten, et al., 2018), which is challenged by the extreme blood flow-induced mechanical forces exerted on the cardiac valves (Hu, et al., 2001; Yalcin, et al., 2011; Kalogirou, et al., 2014; Boselli and Vermot, 2016; Sotiropoulos, et al., 2016; Wissing, et al., 2017).

Most *in vivo* models for valve implantology are large animals, such as pig and sheep, due to the need to surgically insert the implants (Quinn, 2013; Tsang, et al., 2016; Kheradvar, et al., 2017). Despite the translational potential of these animal models given their anatomical and physiological similarities with the human heart, working with large animals has multiple ethical and technical limitations. As an alternative, a mouse model of pulmonary cardiac valve transplantation has recently been developed to study the process of recellularization in tissue-engineered cardiac valves (James, et al., 2015). The introduction of this murine model brought the possibility of using multiple genetic tools. However, the need for surgical implantation of the valve brings two main limitations: i) surgery is particularly challenging due to the small size of the animal; ii) the insertion of a foreign body in the heart generally triggers an exacerbated immune response which may mask the molecular factors underlying the physiological process of valve regeneration. Furthermore, much like the myocardium and other cardiac tissues, mammalian valves display limited regeneration potential (Poss, 2007). Altogether, it becomes particularly challenging to understand which cells are contributing to repopulate the implanted valve leaflets and which molecular factors modulate this process. Thus, it is vital to develop new animal models to study the mechanisms of valve regeneration.

Zebrafish are outstanding in their regenerative capacity (Gemberling, et al., 2013), and multiple studies using this model organism have contributed to uncovering the underlying mechanisms that lead to complex tissue regeneration and to prompt regenerative strategies in mammals. Additionally, the zebrafish cardiac valves share most developmental and morphological features of the mammalian valves (Walsh and Stainier, 2001; Beis, et al., 2005; Scherz, et al., 2008; Martin and Bartman, 2009; Staudt and Stainier, 2012; Pestel, et al., 2016; Steed, et al., 2016; Gunawan, et al., 2019; Schulz, et al., 2019).

Here we used a genetic ablation protocol to decellularize the zebrafish atrio-ventricular (AV) valve and define the regeneration process leading to the formation of new functional valve leaflets. We show that cardiac valve ablation induces the recruitment of endothelial and kidney marrow-derived cells which differentiate into valve cells and secrete extracellular matrix (ECM) to form new cellularized tissue. Interestingly, transcriptome analysis of the ablated and surrounding tissues shows a reactivation of TGFß signaling which promotes cell cycle re-entry. Moreover, by increasing the expression of the ligand Tgfß1b, we were able to enhance new valve cell differentiation, indicating that TGFß acts as a valve pro-regenerative factor with translational potential. Our model provides a new promising approach to identify the cellular and molecular factors regulating new tissue formation during cardiac valve regeneration.

## Results

### Decellularization of the zebrafish atrio-ventricular valve triggers a regenerative program leading to a fully regenerated valve

As in mammals, adult cardiac valves in zebrafish are composed of endothelial cells that surround the extracellular matrix secreted by valve interstitial cells (VICs, Figure 1A-B). Taking advantage of the *TgBAC(nfatc1:Gal4ff)* transgenic line, which expresses Gal4 in cardiac valve cells during development (Pestel, et al., 2016), together with the Nitroreductase/Metronidazole (NTR/Mtz) system (Curado, et al., 2007; Pisharath, et al., 2007), we are able to decellularize the adult AV valve by promoting apoptosis of the VICs (Figure 1D-G). Cell death can be detected by 24 hours post ablation (hpab), and the percentage of surviving cells decreases to below 20% from 72 hpab onward (Figure 1E). Transmission electron microscope (TEM) analyses of uninjured and ablated valve leaflets (Figure 1F-G) show the decellularized ECM, debris of the ablated VICs and rounding of valve endothelial cells (VECs) by 48 hpab. Interestingly, valve decellularization in zebrafish leads to the activation of a regenerative program which results in a newly formed valve with VICs surrounded by ECM (Figure 1H-K). Functional analysis of valve performance by doppler echocardiography in uninjured and at 7 and 14 days post ablation (dpab) shows that decellularization results in valve malfunction and retrograde blood flow in most animals (6/ 8 and 7/12, respectively; Figure S1A-C, Video S1-S3). By 60 dpab, retrograde flow was detected in only 1/9 ablated animals (Figure S1D, D’, Video S4), indicating that the regenerative process leads to the functional recovery of the valve leaflets.

**Figure 1.**
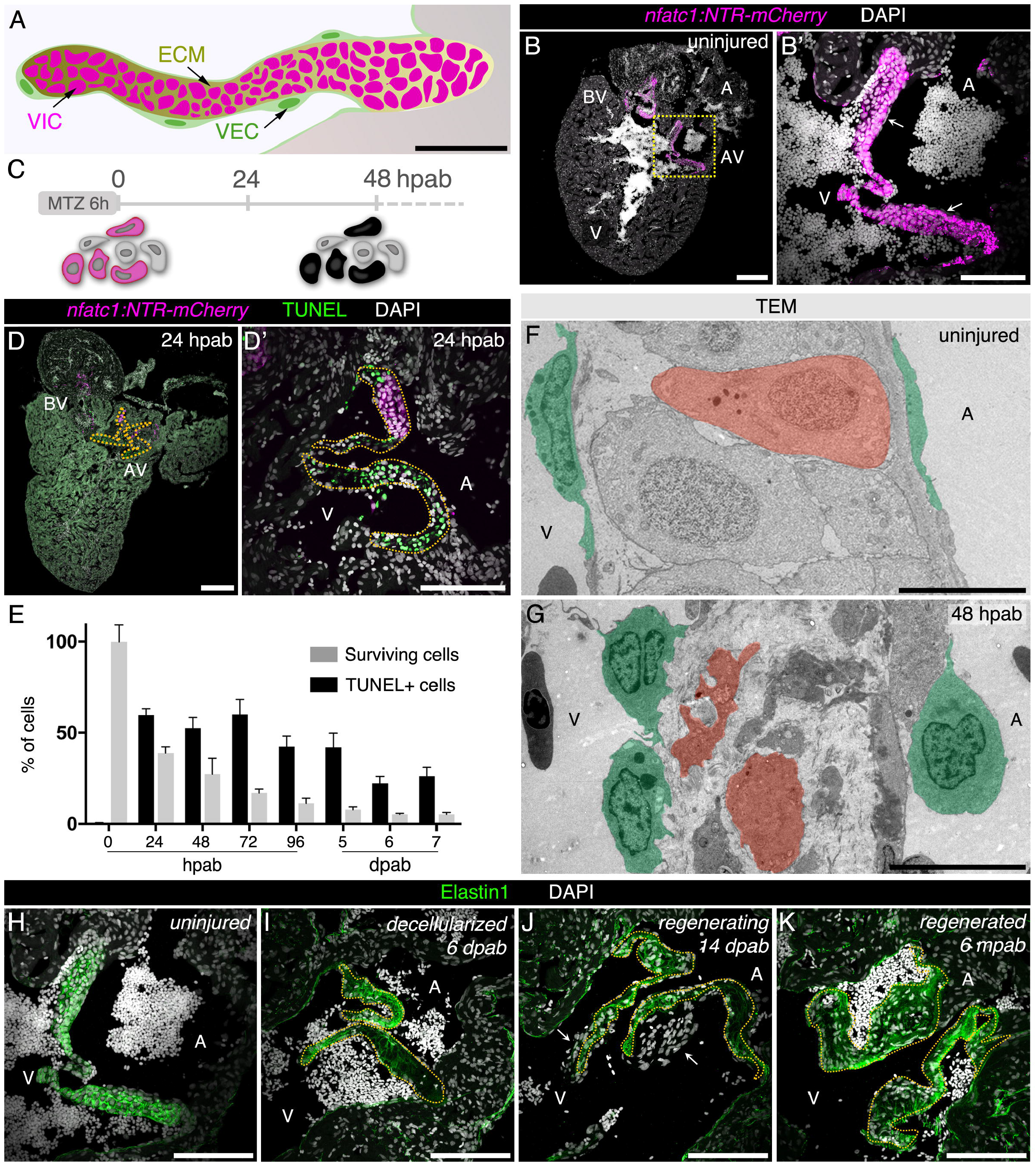
Decellularization of the zebrafish atrio-ventricular valve triggers a regenerative program leading to a fully regenerated valve. (A) Schematics of the zebrafish AV valve leaflet depicting the lining VECs and the ECM-embedded VICs. (B, B’) Cryosection of *Tg(nfatc1:NTR-mCherry)* heart showing NTR-mCherry expression restricted to the VICs in the AV and bulbo-ventricular (BV) valves; NTR-mCherry expression is not detected in the VECs (arrows). Boxed area shown in B’. (C) Ablation protocol using the NTR/Mtz system. TUNEL detection (D, D’) and quantification (E) in ablated hearts shows apoptotic cells only in the valve leaflets. Plot values represent means ± SEM. TEM analyses of uninjured (F) and 48 hpab (G) valve sections show the morphology of the lining VECs (green) and VICs (red) before and after ablation. Representative images of the AV valve regeneration process illustrating the uninjured valve (H), tissue decellularization (I), formation of new valve leaflets (J) and completion of regeneration (K) with ECM labeled by Elastin antibody. Dashed lines delineate the old valve leaflets. A – atrium, V – ventricle. Scale bars: (A) 50 μm, (B, D) 200 μm, (B’, D’, H-K) 100 μm, (F, G) 10 μm.

To understand the overall process of zebrafish cardiac valve regeneration, we analyzed cell cycle re-entry, cell differentiation and new tissue formation (Figure 2). Using EdU labelling to determine cell cycle re-entry, we identified an acute increase in the number of EdU+ cells around the ablated leaflets, peaking at 48-72 hpab (Figure 2A-C). We also detected multiple oval-shaped cells adjacent to the valve leaflets (Figure 2D), a typical morphology of cells re-entering the cell cycle (Théry and Bornens, 2008; Cadart, et al., 2014). Furthermore, we quantified the number of new *nfatc1*+ cells across all time points to determine the extent of valve cell differentiation (Figure 2E-G). New cells accumulated and appeared multi-layered mainly on the ventricular side of the leaflets (Figure 2E-F, H), a process particularly evident between 7 and 21 dpab. The cell differentiation process is accompanied by new ECM secretion, leading to the temporary thickening of the valve leaflets and establishment of a new valve (Figure 2I-N). At 14 dpab, we detected an immature matrix, rich in hyaluronic acid, which subsequently became enriched with Elastin to form a mature, elastic matrix (Figure 2I-L).

**Figure 2.**
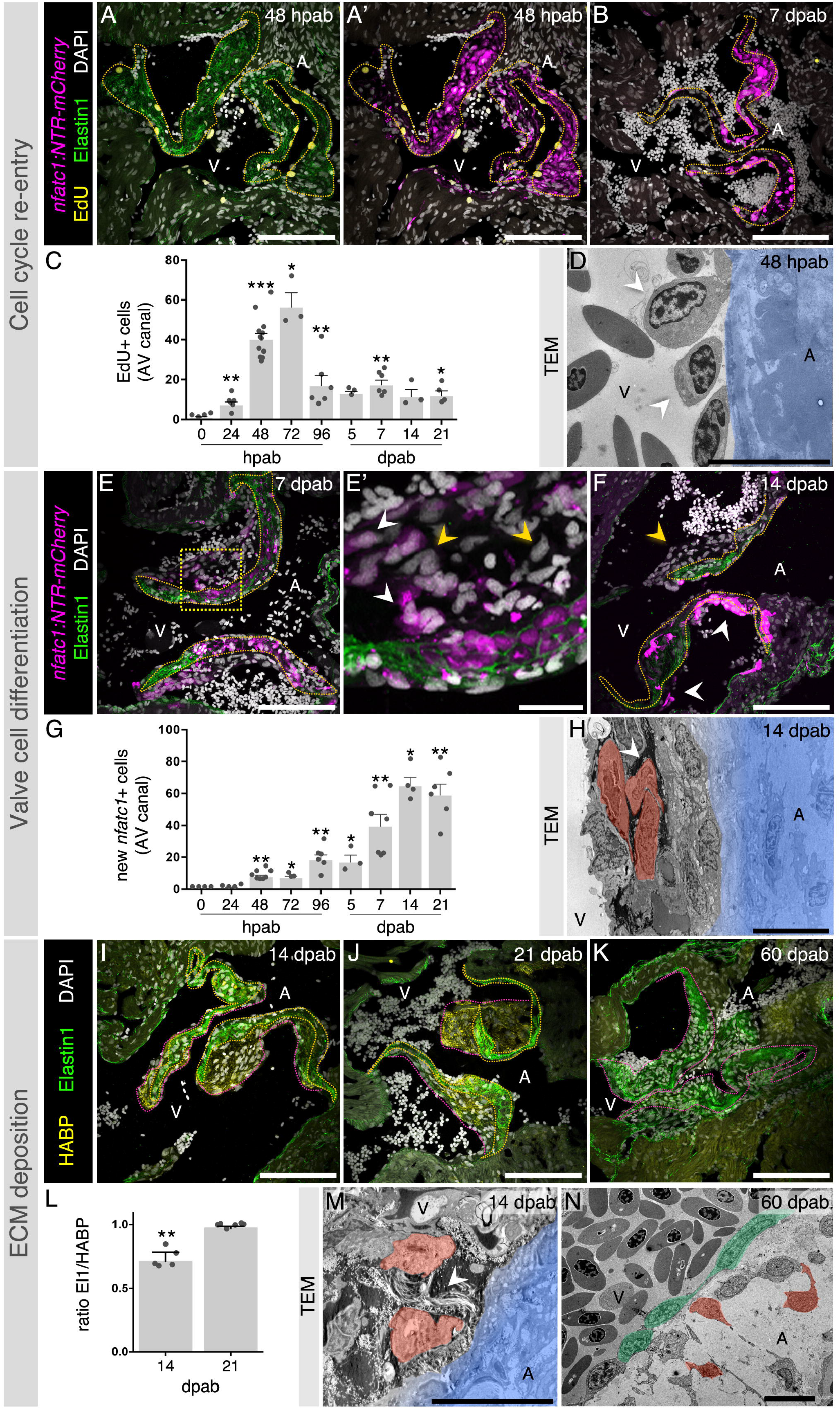
Valve regeneration leads to the formation of new valve leaflets through cell cycle re-entry, new valve cell differentiation and ECM secretion. EdU detection at 48 hpab (A, A’) and 7 dpab (B) and quantification at multiple stages after cell ablation (C). (D) TEM analysis shows oval-shaped cells (arrowheads) adjacent to the decellularized valve (blue). New valve cell differentiation as observed at 7 dpab (E, E’) and 14 dpab (F). Yellow arrowheads point to new *nfatc1*- cells and white arrowheads point to new *nfatc1*+ cells. Boxed area shown in E’. (G) Quantification of new *nfatc1*+ cells in the AV canal. (H) TEM showing new VICs (red) attached to the old matrix (blue) and surrounded by new ECM (arrowhead). Analysis of new ECM deposition at 14 (I), 21 (J) and 60 (K) dpab through detection of Elastin1 and hyaluronic acid (HABP). (L) Quantification of ECM ratios in the valve leaflets at 14 and 21 dpab. TEM images showing VICs (red) secreting ECM (arrowhead) at 14 dpab (M) and fully regenerated valve leaflets at 60 dpab (N; VECs in green). Yellow and pink dashed lines delineate the old and new valve leaflet ECM, respectively. Plot values represent means ± SEM. (\**P*<0.05, \*\**P*<0.01, \*\*\**P* < 0.001 by Mann Whitney test). A – atrium, V – ventricle. Scale bars: (A, A’, B, E, F, I-K) 100 μm, (E’) 20 μm, (D, H, M, N) 10 μm.

### New valve cells differentiate from endothelial and kidney marrow derived-cells

To identify the origin of the new valve cells, we analyzed different endothelial reporter lines and performed cell tracing experiments (Figure 3). Upon analyzing EdU incorporation in various reporter lines at early stages of valve regeneration, we noticed that the majority of the cells re-entering the cell cycle were positive for the*fli1* and *kdrl* driven endothelial reporters (Figure 3A-B’). To quantify the number of EdU+cells exhibiting this endothelial origin, we generated a *Tg(fli1a:H2B-GFP)* line which allows for a prolonged labelling of endothelial cell nuclei (Figure 3B-C). Also, upon staining with a Fli1 antibody, we found that that all the cells exhibiting nuclear Fli1 immunostaining were *kdrl*+ (Figure 3D-D’’), indicating that the Fli1 antibody is a valid endothelial cell marker in our experimental setup.

**Figure 3.**
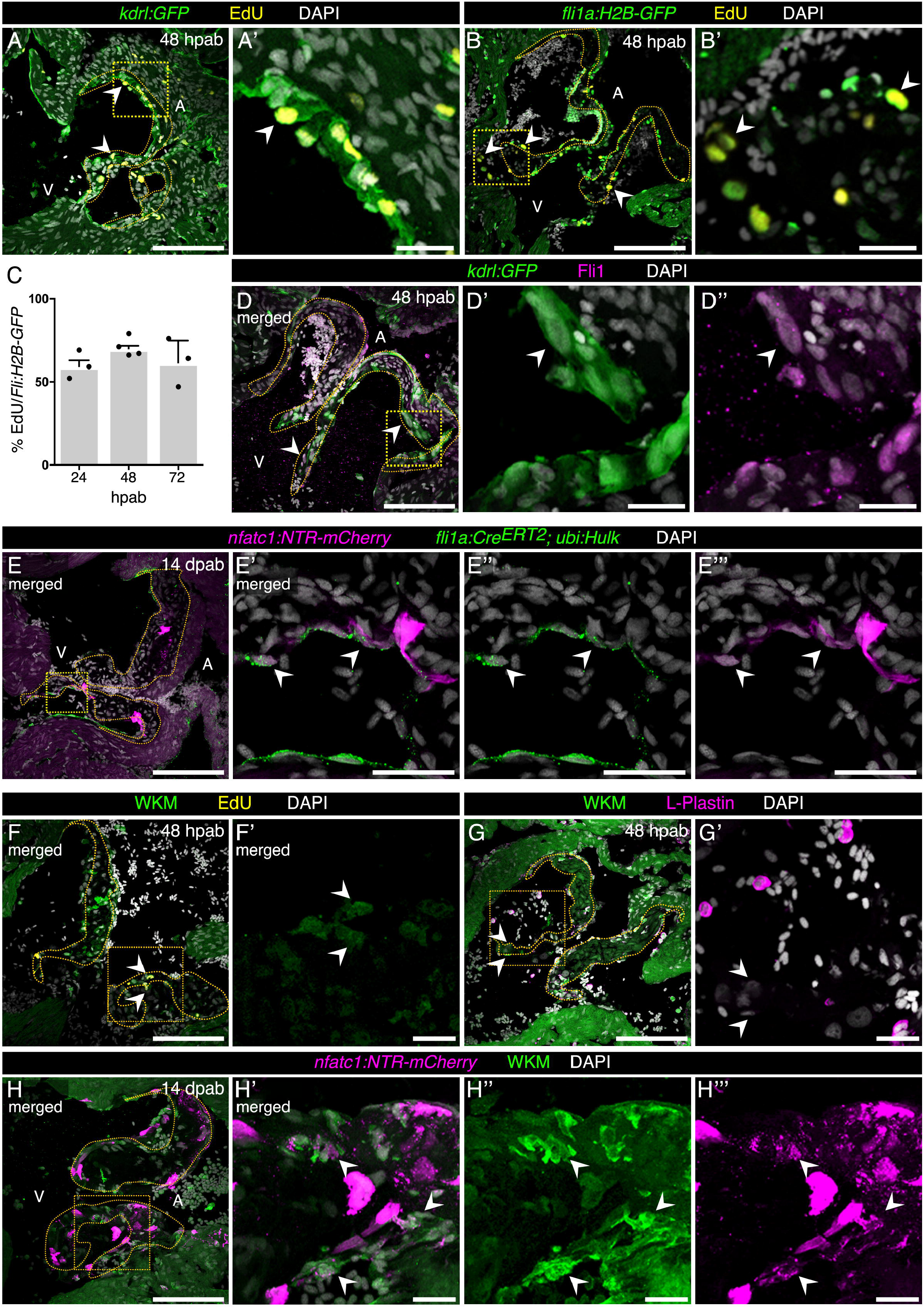
New valve cells differentiate from endothelial and kidney marrow-derived cells. Characterization of EdU co-localization with endothelial reporters (arrowheads) at 48 hpab including *kdrl:GFP* (A, A’) and*flia:H2B-GFP* (B, B’). (C) Quantification of the percentage of EdU+ cells positive for *flia:H2B-GFP.* Plot values represent means ± SEM. (D) Co-localization of *kdrl* reporter with Fli1 antibody (arrowheads). Boxed areas shown in D’ and D’’. (E-E’’’) Cre/lox cell tracing experiments showing contribution of endothelial cells to *nfatc1*+ cells (arrowheads) at 14 dpab (n=6). Boxed areas shown in E’-E’’’. WKM-derived cells positive for EdU (F-F’) and negative for the immune cell marker L-Plastin (G-G’) at 48 hpab. Boxed areas shown in F’ and G’. (H-H’’’) Contribution of circulating WKM-derived cells to new *nfatc1*+ cells (arrowheads) at 14 dpab (n=4). Boxed areas shown in H’-H’’’. Dashed lines delineate the valve leaflets. A – atrium, V – ventricle. Scale bars: (A, B, D, E, F, G, H) 100 μm, (A’, B’, D’-D’’, E’- E’’’, F’, G’, H’-H’’’) 20 μm.

To test whether endothelial cells were differentiating to VICs, we took advantage of the Cre/lox system (Sánchez-Iranzo, et al., 2018) and induced reporter recombination in endothelial cells prior valve cell ablation. With this approach, we saw that an average of 10±6 *fli1*+ endothelial cells (per section) which recombined to express GFP prior to valve cell ablation became *nfatc1*+ by 14 dpab (Figure 3E-E’’’). Therefore, we were able to confirm that endothelial cells differentiate to give rise to the new valve cells (Figure 3E-E’’’). Additionally, we determined the contribution of circulating cells from hematopoietic tissue to cardiac valve regeneration using whole kidney marrow (WKM) as previously described (Traver, et al., 2003). To this end, we transplanted GFP-labelled WKM from a *Tg(ubi:switch);Tg(nfatc1:NTR-mCherry)* fish into irradiated *Tg(nfatc1:NTR-mCherry)* adult animals prior to valve cell ablation. We found that at 48 hpab some of the EdU+ cells in the AV canal were derived from WKM (Figure 3F-F’) and that, although the majority (86,3±0.2%) of the WKM-derived cells were positive for immune cell markers, some of them were negative (Figure 3G-G’), suggesting that they might be valve cell precursors. Interestingly at 14 dpab we identified that an average of 6±2 (per section) of the *nfatc1*+ cells also derived from WKM (Figure 3H-H’’’). Altogether, these data show that endothelial as well as circulating hematopoietic cells differentiate to new valve cells.

### TGFß signaling enhancement increases cell cycle re-entry and differentiation of new *nfatc1*+ cells during AV valve regeneration

Once we had defined the main stages of AV valve regeneration, we performed a transcriptome analysis to identify molecular signatures associated with the regenerative process. For this study, we had to take into consideration that the AV valve leaflets are in close proximity to multiple tissues, including the epicardium, myocardium and endocardium. Additionally, multiple cell types can easily access the valve leaflets, including blood, immune and other circulating cells. These points make it particularly difficult to select reporter lines to sort all the possible cell types contributing to zebrafish valve regeneration. Furthermore, the use of dissociation protocols could potentially mask processes, such as endothelial-to-mesenchymal transition (EndoMT) or cell-ECM interactions, which we expected to be relevant for cardiac valve regeneration. Therefore, we decided to use laser capture microdissection (LCM; (Datta, et al., 2015)) to isolate uninjured, 48 hpab and 21 dpab valve leaflets and surrounding tissue (Figure S2A) which included epicardium, myocardium, endocardium as well as circulating cells (Figure S2B). Principal component analysis (PCA; Figure S2C) shows that the 21 dpab samples are more similar to an uninjured condition than to the 48 hpab samples. Gene set enrichment analysis of differentially expressed genes during the regenerative process (Figure S2D) indicate a role the immune system and cell cycle, as well as ECM organization. By selecting VIC specific markers, we observed a clear downregulation upon decellularization at 48 hpab, and a relative upregulation at 21 dpab during new valve cell redifferentiation (Figure S2E). Analysis of the 48 hpab samples showed a differential expression of 2115 genes, from which 951 appeared to be upregulated (Figure S2F). In order to identify secreted proteins potentially recruiting new cells to regenerate the ablated AV valve, we compared the list of upregulated genes with the secreted factor genes from the zebrafish matrisome ((Nauroy, et al., 2018); Figure S2G). We identified a total of 12 genes, including *tgfß1b,* which has previously been described to play a critical role in valve development, specifically in the regulation of EndoMT, as well as in VIC proliferation and differentiation (Mercado-Pimentel and Runyan, 2007; Conway, et al., 2011).

Taking advantage of the highly sensitive RNAscope-based *in situ* hybridization, we were able to detect *tgfβ1b* transcripts in cells surrounding the ablated leaflets at 48 hpab (Figure S3A-B’’). When co-staining for endothelial and immune cell markers, we identified that the majority of the cells expressing *tgfβ1b* were endothelial cells (Figure S3C, D). To further determine the activation of the signaling pathway, we immunostained for pSmad3, a downstream effector of TGFß signaling (Derynck and Budi, 2019) in combination with an endothelial marker (Figure S3E, E’) and EdU (Figure S3F, F’). We observed that most pSmad3+ cells were Fli1+, and that most of the EdU+ cells were pSmad3+, suggesting TGFß signaling activation in cells potentially contributing to valve regeneration. Therefore, we performed loss- and gain-of-function experiments of TGFß signaling during cardiac valve regeneration to address its role in cell cycle re-entry and new valve cell differentiation (Figure 4). We used the previously validated TGFß inhibitor SB431542 (Jazwinska, et al., 2007; Chablais and Jazwinska, 2012), and generated a heat-shock inducible dominant negative form of the zebrafish Tgfß receptor I, Alk5a (Figure 4A, B). Additionally, we developed a *nfatc1* driven *tgfβ1b* overexpression transgenic line. To validate the inhibition and activation tools, we analyzed the number of pSmad3+ cells around the ablated leaflets at 48 hpab (Figure S4). We confirmed that SB431542 treatment and *tgfβ1b* overexpression induced a reduction and increase in the number of pSmad3+ cells, respectively (Figure S4A-E). In fact, *tgfβ1b* overexpression (Figure S4D-F) nearly doubled the number of pSmad3+ cells at 48 hpab in comparison to control. For *DNalk5a* overexpression (Figure S4C, E, H), we submitted the ablated fish to a single mild heat-shock at 20 hpab (Figure 4A). Quantification of the number of pSmad3+ cells at 48 hpab showed no clear reduction 28 hours after heat shock. Therefore, we validated the effectiveness of this dominant negative protein *in vitro* (Figure S4H). While the zebrafish wild-type protein further increased the endogenous SBE motif-dependent luciferase expression upon TGFß treatment, the dominant negative protein abrogated this response. In parallel, we used as an additional control the previously validated human dominant negative Alk5a (Wieser, et al., 1995) which, like zebrafish DNAlk5a, caused a loss in the response to TGFß. Once these tools were validated, we determined whether TGFß signaling was required for cell cycle re-entry at 48 hpab (Figure 4). Notably, we observed that pharmacological inhibition of TGFß signaling and overexpression of *DNalk5a* led to a reduction in the number of EdU+ cells by almost 30 and 50%, respectively (Figure 4C-F). Importantly, *nfatc1*-driven overexpression of the *tgfβ1b* ligand increased the number of EdU+ cells at 48 hpab by 50% (Figure 4C, G). It also doubled the number of *nfatc1+* cells at 14 dpab (Figure 4H-J), thus promoting valve cell differentiation. Altogether, these data indicate that TGFß signaling promotes valve regeneration.

**Figure 4.**
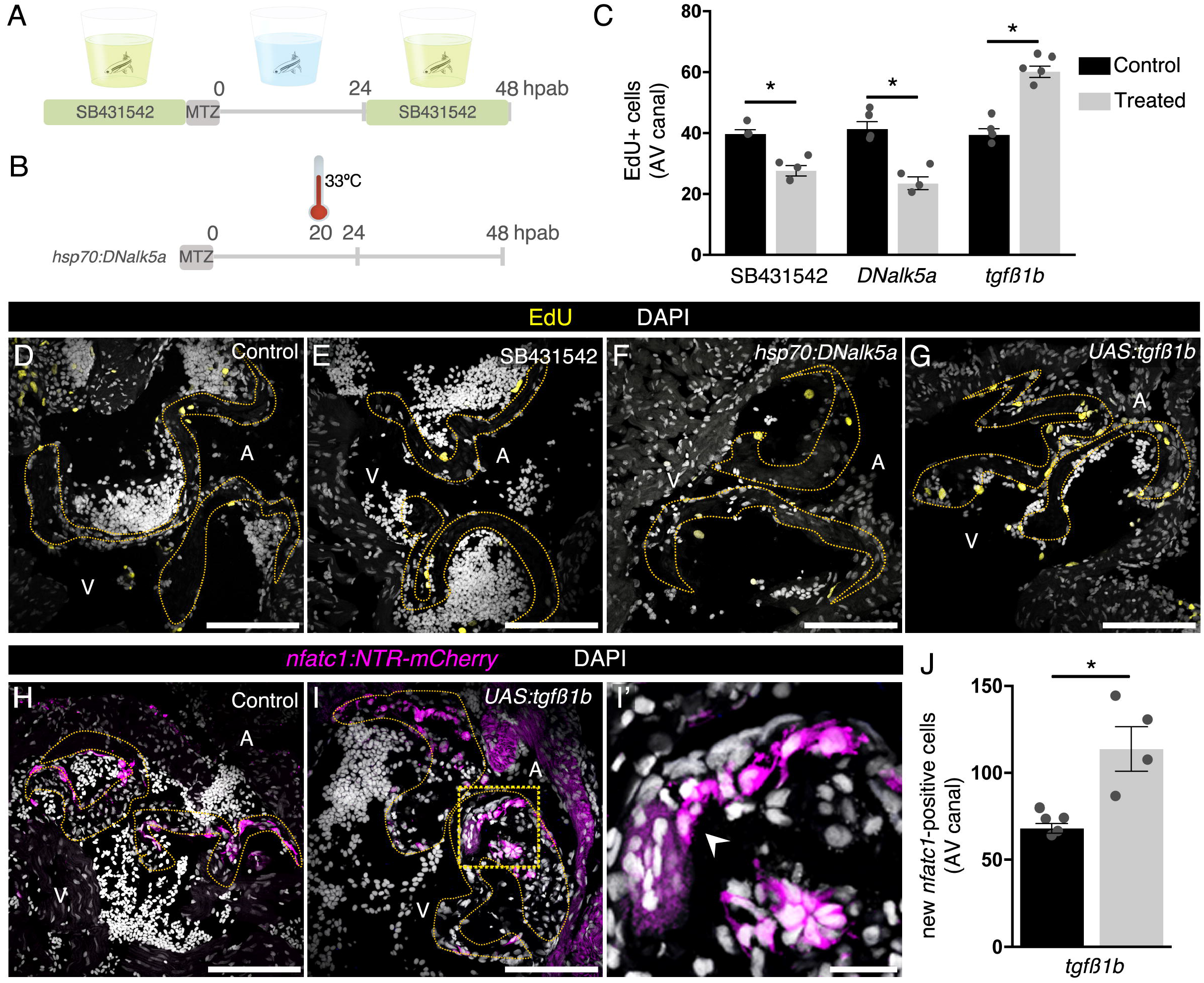
Manipulation of TGFß signaling alters cell cycle re-entry and differentiation of *nfatc1*+ cells during cardiac valve regeneration. Schematics of the drug (A) and heat-shock (B) treatments. EdU quantification (C) and detection at 48 hpab in control samples (D), after SB431542 treatment (E) and overexpression of *DNalk5a* (F) or *tgfβ1b* (G). Characterization of new valve cell differentiation at 14 dpab in control (H) and upon continued expression of *tgfβ1b* (I, I’). Arrowhead points to new *nfatc1*+ cells. Boxed area shown in I’. (J) Quantification of new *nfatc1*+ valve cells at 14 dpab. Plot values represent means ± SEM. (\**P* <0.05 by Mann Whitney test). Dashed lines delineate the AV valve leaflets. A – atrium, V – ventricle. Scale bars: (D-I) 100 μm, (I’) 20 μm. See also Figure S4.

## Discussion

Since the first description of an artificial valve over 60 years ago (Hufnagel, et al., 1954), surgeons have been moving from the use of mechanical to living cardiac valve prostheses which require recellularization of the matrix and tissue growth (De Visscher, et al., 2010; Weber, et al., 2013; Driessen-Mol, et al., 2014; Tudorache, et al., 2016; Reimer, et al., 2017; Emmert, et al., 2018; Baldwin and Tolis, 2019). By developing the first model of adult cardiac valve regeneration in the highly regenerative zebrafish, we provide new insights into the processes leading to functional re-establishment of the decellularized AV valve (Figure S5), including the origin of new valve cells, the genes involved in recruiting them, and the *de novo* tissue formation.

### Multi-cellular contribution to cardiac valve regeneration

Multiple cell types have been proposed as VIC precursors, including ECs, myofibroblasts, bone marrow-, peripheral blood-, and adipose-derived stem cells (Schenke-Layland, et al., 2003; Sutherland, et al., 2005; Dohmen, et al., 2006; Schmidt, et al., 2010; Harrington, et al., 2011; Kennamer, et al., 2016). However, most of these cells have never been shown to contribute to valve development or recellularization *in vivo* except for ECs, the main VIC source in amniotes (Snarr, et al., 2008; Jain, et al., 2011; Wessels, et al., 2012; MacGrogan, et al., 2014; Liu, et al., 2018). We show that upon injury, the endothelium surrounding the leaflets re-enters the cell cycle and acquires a morphology resembling that previously described in highly proliferative embryonic and postnatal mouse valves (Anstine, et al., 2016). Therefore, valve ablation induces endothelial activation similar to what happens during development, resulting in new valve cell differentiation. We also show that cells deriving from WKM, the fish counterpart to the mammalian bone marrow (Traver, et al., 2003), contribute to the formation of new valve cells. This observation is in agreement with previous reports suggesting that the majority of cells repopulating a tissue-engineered valve implant in lamb are blood-derived (Reimer, et al., 2017) and that hematopoietic stem cells (HSC) differentiate to VICs in non regenerating adult mice valves (Visconti, et al., 2006). Therefore, we show the first *in vivo* evidence that ECs and WKM-derived cells contribute to cardiac valve regeneration. Moreover, we propose that new valve cell differentiation and *de novo* tissue formation is orchestrated by multiple cell types.

### Identifying molecular regulators of new valve cell recruitment

Recent studies have focused on identifying and supplementing valve scaffolds with molecular factors promoting valve cell recruitment (Jordan, et al., 2012; Jana, et al., 2014). As in other regeneration models, we hypothesized that valve regeneration would rely on the reactivation of developmental signaling cues such as BMP, Notch and TGFß pathways (Yamagishi, et al., 2009; Conway, et al., 2011; de la Pompa and Epstein, 2012; Kruithof, et al., 2012). By combining a sophisticated approach to microdissect the regenerating tissues with transcriptomic analysis, we identified multiple genes potentially associated with early stages of valve regeneration, including the secreted TGFß ligand *tgfβ1b.* Despite the fact that TGFß signaling deregulation in homeostasis induces pathological phenotypes in cardiac valves, such as ECM calcification (Anderton, et al., 2011; Wirrig and Yutzey, 2014; White, et al., 2015; Dutta and Lincoln, 2018), previous groups suggested that it may have a role in tissue regeneration, EndoMT and valve recellularization (Jazwinska, et al., 2007; Liu and Gotlieb, 2008; Benton, et al., 2009; Deng, et al., 2011; Li and Gotlieb, 2011; Chablais and Jazwinska, 2012). Using multiple genetic tools, we show that TGFß signaling is required at early regeneration stages and its enhancement promotes cell cycle re-entry and new valve cell differentiation. Interestingly, ECs were the main source of the Tgfß1b ligand, as well as the main responders with TGFß signaling activation via Smad3. Therefore, we propose that TGFß signaling regulates EC proliferation and EndoMT in AV valve regeneration in an autocrine manner, promoting VIC differentiation. These data suggest that the regenerative process can be enhanced by manipulation of single molecules, therefore providing new therapeutic targets in cardiac valve replacement.

### Formation of new valve tissue

As opposed to what was initially expected, despite the presence of immune cells within the decellularized ECM (data not shown), the new valve cells do not seem to invade the old matrix. Instead, newly differentiated valve cells use the decellularized matrix as a scaffold to adhere on the ventricular surface of the leaflets and build a new valve. These cells have the ability to secrete ECM components also present in native leaflets, conferring stiffness and the elastic properties needed for the functional recovery of the AV valve. Overall, the ultimate goal in cardiac valve replacement is to promote endogenous *de novo* tissue formation. This new tissue ensures growth and maintenance of the biomechanical properties of the valve leaflets particularly relevant for children and young adults due to somatic growth (Driessen-Mol, et al., 2014; Cheung, et al., 2015; David, 2016; Reimer, et al., 2017). Recently biodegradable synthetic matrices have been developed towards promoting new tissue formation and growth of living valves (Bouten, et al., 2012; Kluin, et al., 2017). Since mammalian models display limited regeneration rates, we propose the zebrafish model as a platform to determine the mechanisms underlying the intrinsic ability to form new tissue and grow new valve leaflets.

## Supporting information

Echocardiography uninjured

Echocardiography 7 dpab

Echocardiography 14 dpab

Echocardiography 60 dpab

## Acknowledgments

We would like to thank Jill de Jong and Sean McConnell for insights regarding WKM transplantation; Matteo Perino for the p3TP:Luc2, SV40:hRLuc2 plasmid and Paul Martin for the L-plastin antibody; Simon Howard, Radhan Ramadass and all the fish facility staff for technical support; Christian Helker for discussions; João Cardeira da Silva, Inês Cristo, Felix Gunawan, Carol Yang and Honorine Destain for critical comments on the manuscript. Research in the Stainier lab is supported in part by the Max Planck Society and the European Union (ERC).

## Author Contributions

Conceptualization, A.B.B. and D.Y.R.S.; Methodology, A.B.B., S.R., G.L.M.B., S.G., C.K., H.S.I., D.I., J.P. and D.Y.R.S.; Validation, A.B.B., S.R. and G.L.M.B.; Formal Analysis, A.B.B., S.R., G.L.M.B., S.G. and C.K.; Investigation, A.B.B., S.R., G.L.M.B., S.G., J.P., Resources, H.S.I., D.I., S.P., N.M., D.B. and D.Y.R.S.; Writing – Original Draft, A.B.B. and D.Y.R.S.; Writing – Reviewing & Editing, all; Visualization – A.B.B., Supervision, A.B.B. and D.Y.R.S.; Project Administration, A.B.B. and D.Y.R.S.; Funding Acquisition, D.Y.R.S.

## Declaration of Interests

The authors declare no competing interests.

---

**Figure S1.**
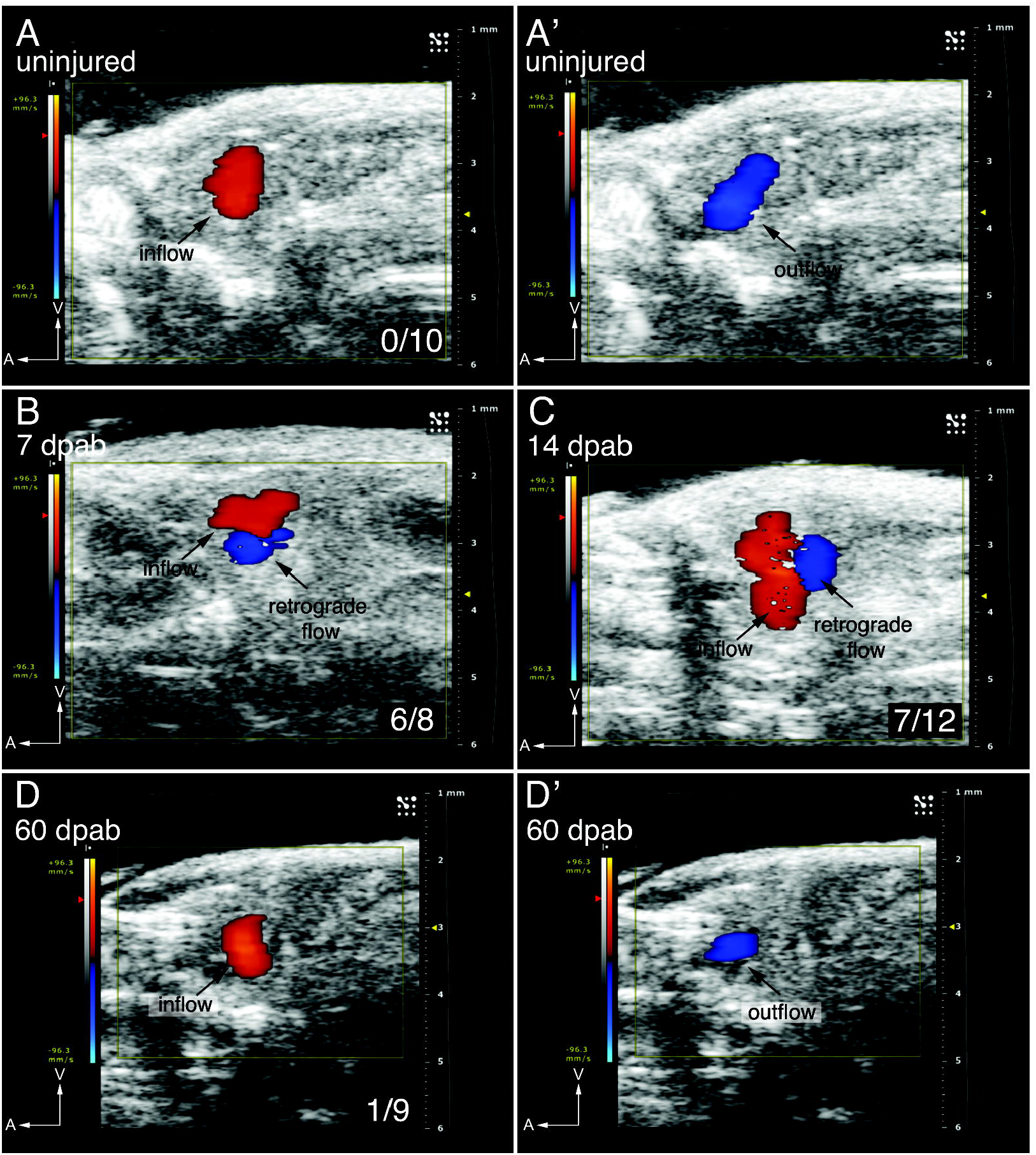
Regenerated valves recover from functional impairment induced by the decellularization protocol. Doppler echocardiography analysis of cardiac function in an uninjured fish (A, A’) showing unperturbed blood inflow (red) and outflow (blue) without signs of regurgitation (see also Video S1). Retrograde blood flow is detected in 6/8 animals at 7 dpab (B; see also Video S2) and 7/12 animals at 14 dpab (C; see also Video S3). (D, D’) Clear separation of incoming and outgoing blood in the same specimen at 60 dpab, as observed in 8/9 of the ablated animals (see also Video S4). A – anterior, V – Ventral.

**Figure S2.**
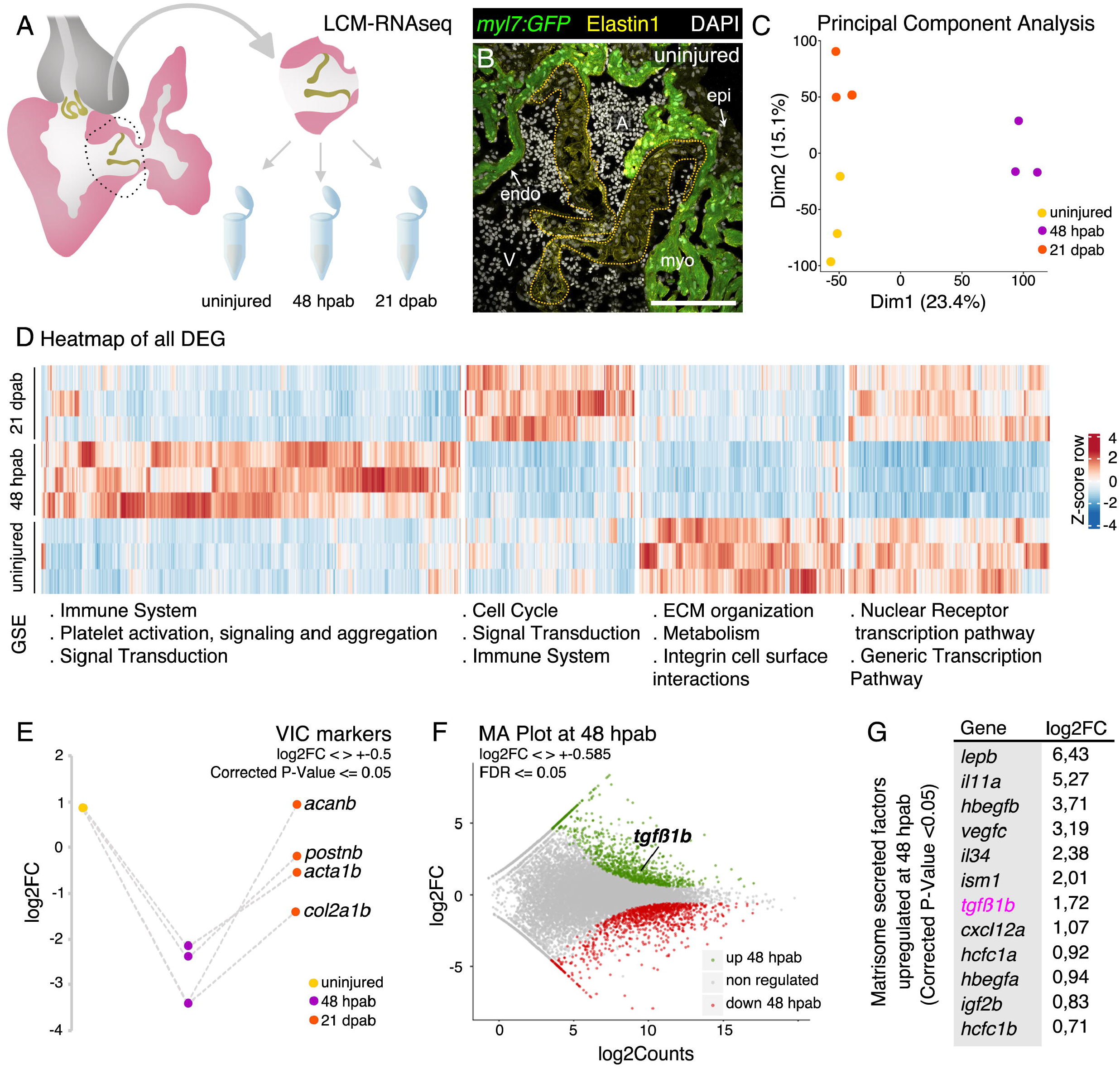
LCM-RNAseq transcriptome analysis. (A) Scheme of the AV tissues isolated through LCM for RNAseq analysis. (B) Cryosection of *myl7:GFP* heart showing the endocardium (endo), myocardium (myo) and epicardium (epi) in the vicinity of the AV valve leaflets. (C) PCA analysis of RNAseq samples based on DESeq normalized and rlog transformed counts of all genes. (D) Heatmap with cluster and gene set enrichment (GSE) analyses of DEGs with main biological processes regulated in each cluster. (E) Regulation of VIC markers relative to uninjured samples. MA plot from DEGs (F) and list of matrisome secreted factors upregulated at 48 hpab (G), including Tgfß1b. A – atrium, V – ventricle. Scale bar: (B) 100 μm.

**Figure S3.**
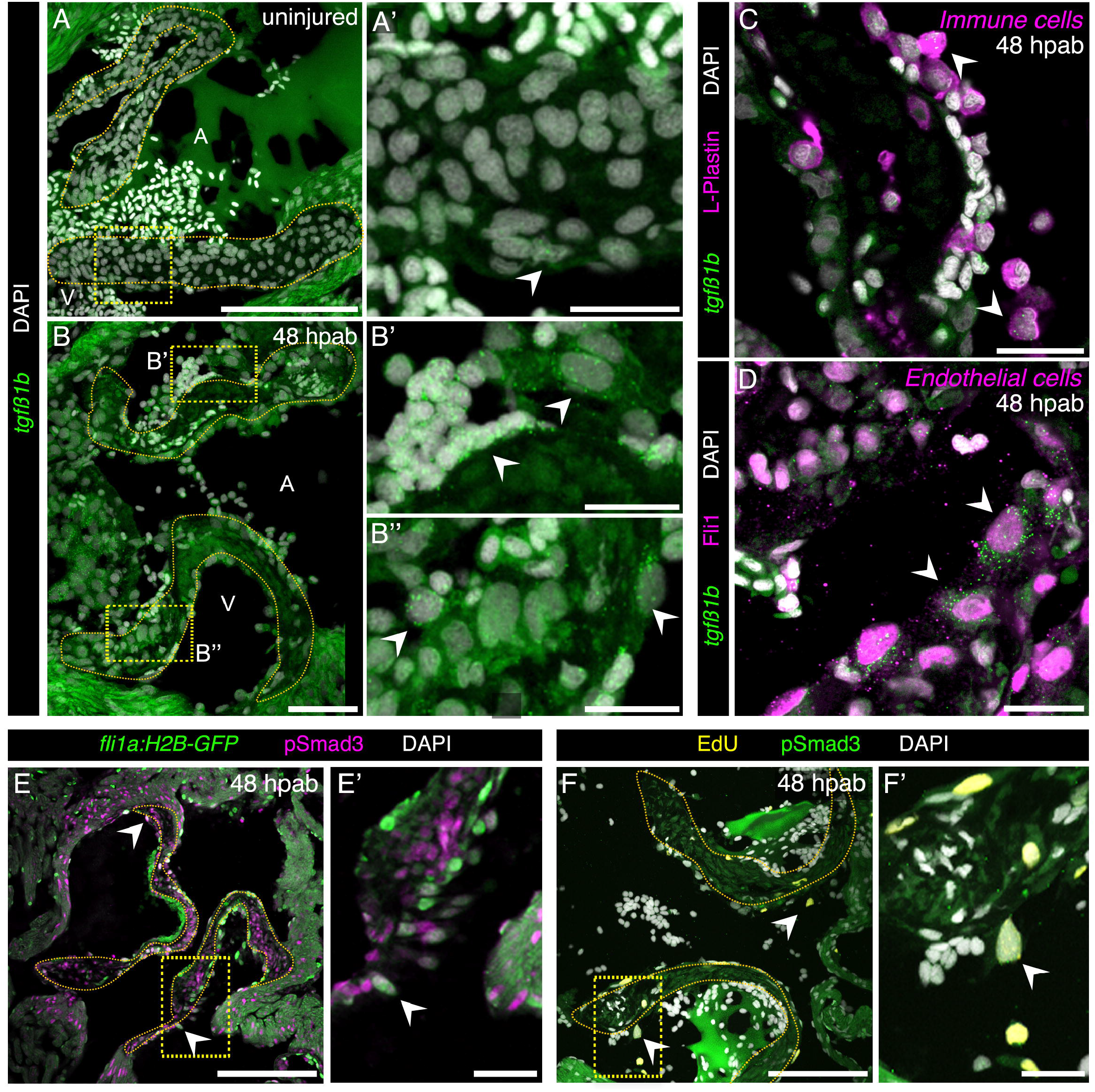
Cell cycle re-entry is accompanied by an increase in *tgfβ1b* expression and activation of TGFß signaling in endothelial cells. RNAscope *in situ* hybridization of *tgfβ1b* (arrowheads) in heart cryosections in uninjured (A, A’) and 48 hpab (B, B’, B’’) AV valves. Boxed areas shown in A’, B’ and B’’. *In situ* hybridization for *tgfβ1b* (arrowheads) co-stained for immune (C) and endothelial cell markers (D). Characterization of TGFß signaling activation by pSmad3 immunodetection (arrowheads) together with an endothelial reporter (E, E’) and EdU (F, F’). Boxed areas shown in E’ and F’. Dashed lines delineate the AV valve leaflets. A – atrium, V – ventricle. Scale bars: (A, B, E, F) 100 μm, (A’, B’, B’’, C, D, E’, F’) 20 μm.

**Figure S4.**
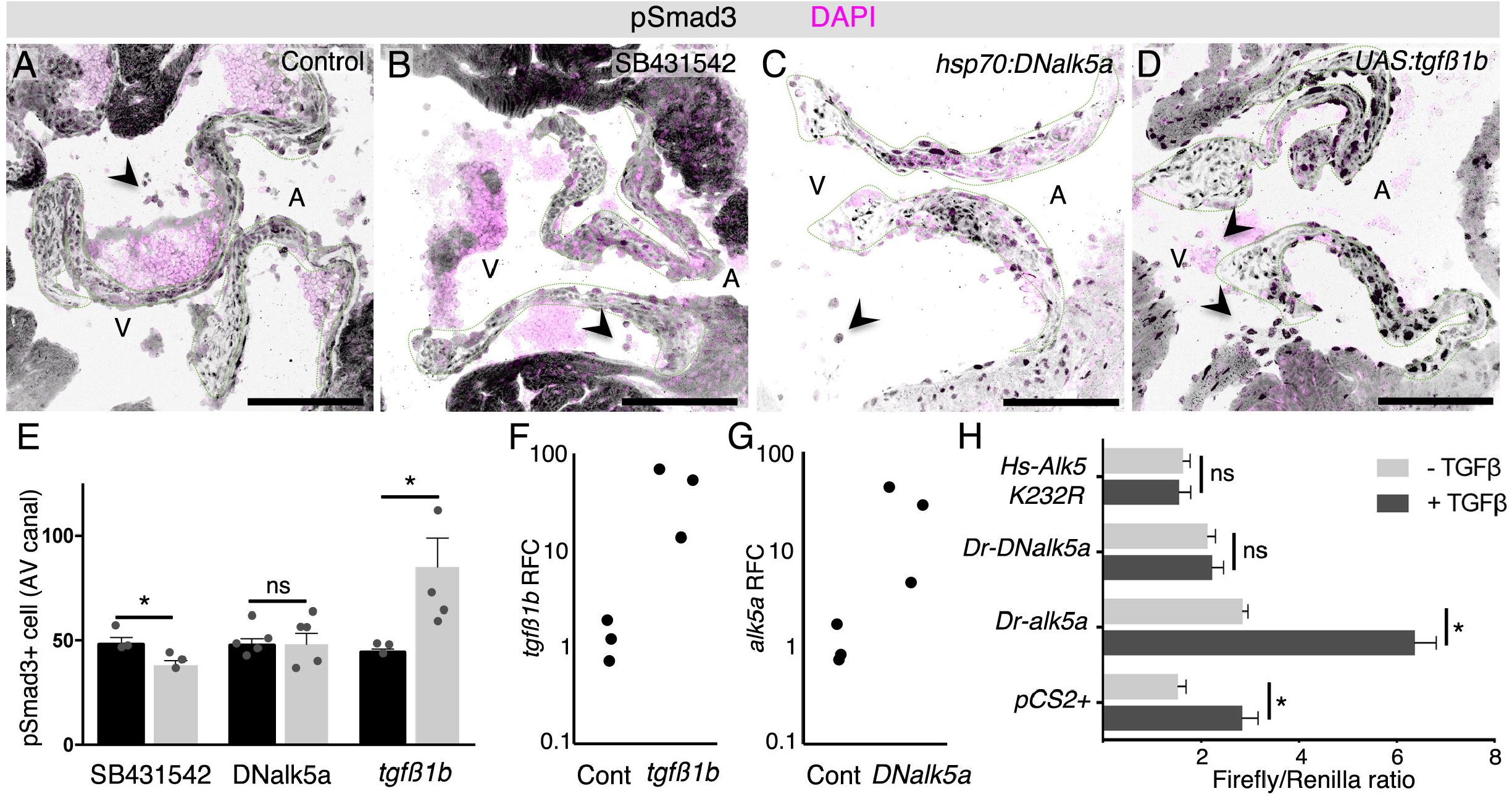
Validation of the TGFß signaling modulation tools. pSmad3 detection at 48 hpab to determine levels of TGFß signaling activation in controls (A), after SB431542 treatment (B) and overexpression of *DNalk5a* (C) or *tgfβ1b* (D). (E) Quantification of pSmad3+ cells in the AV canal. qPCR analysis of mRNA relative fold change (RFC) of *tgfβ1b* (F) and *alk5a* (G) in the overexpression lines. (H) *In vitro* luciferase assay of zebrafish Alk5a wild-type and dominant negative proteins, and human ALK5 dominant negative protein in the absence/presence of the TGFß ligand. Plot values represent means ± SEM. (ns – non-significant, \**P*<0.05 by Mann Whitney test). Dashed lines delineate the AV valve leaflets. A – atrium, V – ventricle. Scale bars: 100 μm.

**Figure S5.**
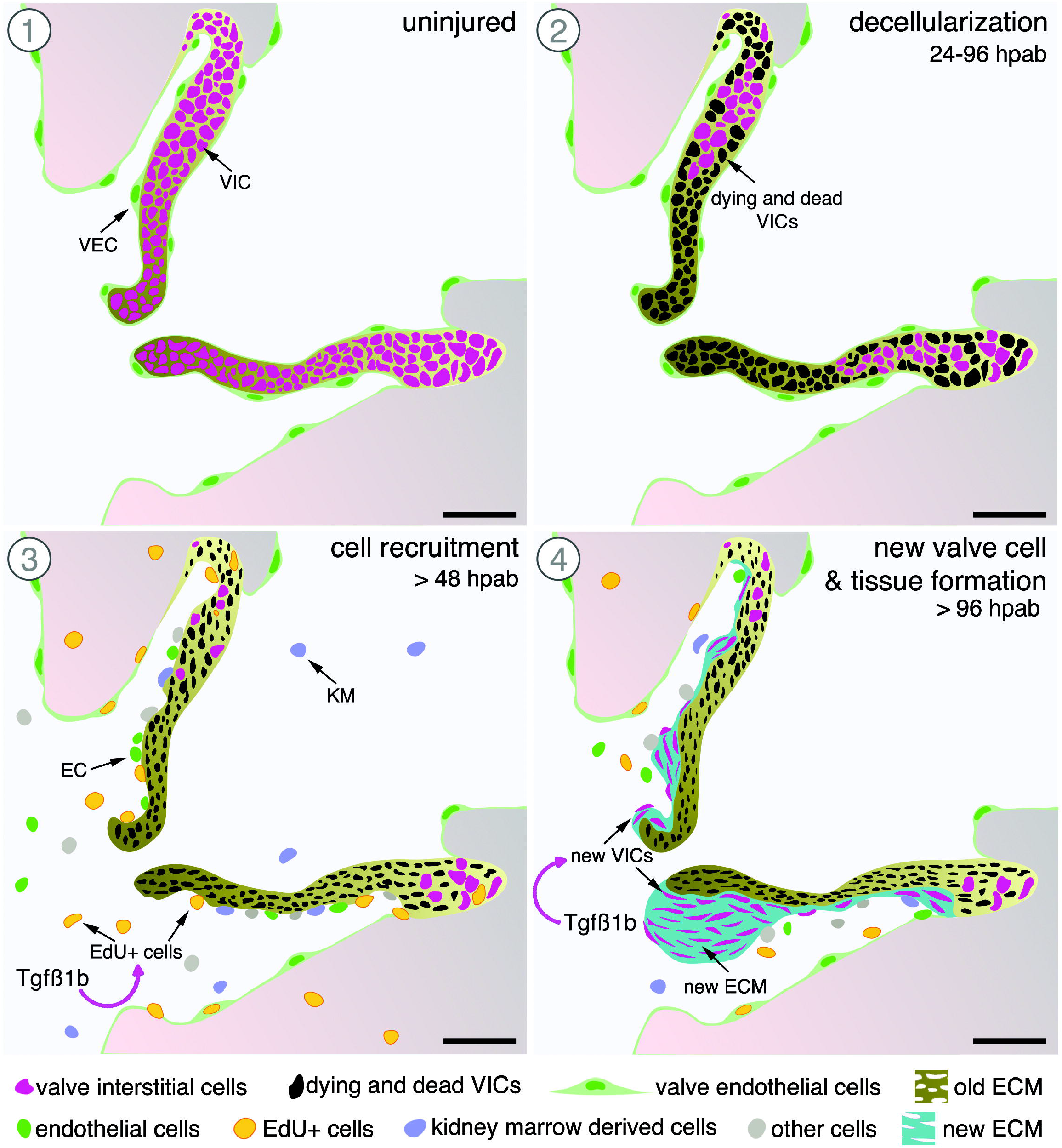
Zebrafish cardiac valve regeneration model. AV valve ablation leads to the recruitment of endothelial and kidney marrow cells, which will contribute to the formation of new valve leaflets. TGFß signaling promotes valve regeneration by regulating cell cycle re-entry and consequent valve cell differentiation which leads to new ECM secretion. VIC – valve interstitial cell, VEC – valve endothelial cell, EC – endothelial cell, KM – kidney marrow. Scale bars: 50 μm.

Video S1 - Doppler echocardiography analysis of cardiac function in an uninjured fish showing unperturbed blood inflow (red) and outflow (blue) without signs of regurgitation (Related to Figure S1A-A’).

Video S2 - Doppler echocardiography analysis at 7 dpab showing retrograde blood (Related to Figure S1B).

Video S3 - Doppler echocardiography analysis at 14 dpab showing retrograde blood (Related to Figure S1C).

Video S4 - Doppler echocardiography analysis at 60 dpab showing a clear separation of incoming and outgoing blood (Related to Figure S1D-D’).

## STAR Methods

### LEAD CONTACT AND MATERIALS AVAILABILITY

Further information and requests for resources and reagents should be directed to and will be fulfilled by the Lead Contact, Didier Y. R. Stainier.

### EXPERIMENTAL MODEL AND SUBJECT DETAILS

#### Zebrafish and cell lines

All zebrafish husbandry was performed under standard conditions, according all institutional (MPG) and national ethical and animal welfare guidelines.

Wild-type or transgenic male and female zebrafish used in this study were derived from the AB strain. Animals were used for ablations from 3 to 6 months post fertilization (mpf). Rearing system and incubators were maintained at 28°C with a light cycle of 10 hours dark/14 hours light. All transgenic animals were analyzed as hemizygotes except for the *UAS:tgfβ1b* and the *hsp70:DNalk5a* strains.

Luciferase assays were performed using HEK-293T cells.

### METHOD DETAILS

#### Transgenic lines and plasmids

The following lines were used:

*TgBAC(nfatc1:Gal4ff)^mu286^* (Pestel, et al., 2016) together with *Tg(UAS:NTR-mCherry)^c264^* (Davison, et al., 2007), abbreviated *nfatc1:NTR-mCherry; Tg(kdrl:NLS-EGFP)^zf109^* (Blum, et al., 2008), abbreviated *kdrl:GFP; Tg(fli1a:Cre-ERT2)^cn9^* (Sánchez-Iranzo, et al., 2018), abbreviated *fli1a:Cre^ERT2^; Tg(−3.5ubb:LOXP-LacZ-LOXP-eGFP)^cn2^* (Di Donato, et al., 2016), abbreviated Hulk; *Tg(−3.5ubb:loxP-EGFP-loxP-mCherry)^cz1701^* (Mosimann, et al., 2011), abbreviated *Tg(ubi:Switch); Tg(myl7:GFP)^f1^* (Burns, et al., 2005), abbreviated *myl7:GFP.*

To generate *Tg(fli1a:H2B-GFP)^bns319^*, abbreviated *fli1a:H2B-GFP,* a 2kb *fli1a* regulatory region was cloned upstream of H2B-GFP sequence. The *fli1a* promoter sequence was amplified using the following primers: forward 5’-TTGGAGATCTCATCTTTGAC-3’ and reverse 5’ - GGTGGCGCTAGCTTCGCGTCTGAATTAATTCC-3’.

To generate *Tg(5xUAS-hsp70:tgfβ1b-p2A-H2B-GFP)^bns315^,* abbreviated *UAS:tgfβ1b, tgfβ1b* (XM_687246.8) was cloned downstream of the *5xUAS-hsp70* regulatory sequence. The full-length *tgfβ1b* coding sequence was amplified using the following primers: forward 5’-ATGAAGGCGGAGAGTTTATT-3’ and reverse 5’- CAACACTTGCAGGTTTTCAC-3’.

To generate *Tg(hsp70:DNalk5a-p2A-H2B-GFP)^bns317^,* abbreviated *hsp70:DNalk5a,* a partial coding sequence of *alk5a* (NM_001037683.2) lacking the intracellular STK (serine-threonine kinase) domain was cloned downstream of the *hsp70* promoter sequence. The *DNalk5a* coding sequence was amplified using the following primers: forward 5’- ATGAGCTCCGCCGCGATGG-3’ and reverse 5’- GGTCCTGGCGATGGTCCGTT-3’.

All vectors had *Tol2* elements to facilitate genome integration.

#### Luciferase assay and plasmids

For the luciferase expression we generated p3TP:Luc2, SV40:hRLuc2 plasmid from the pGl4.14 GLO backbone (Promega), by inserting the promoter from p3TP-Lux (Addgene) upstream of the luciferase (Luc2) and the human Renilla luciferase coding sequence (amplified from pGl4.79, Promega) to serve as an internal control. The 3TP promoter sequence was amplified using the following primers: forward 5’-CAGCTGAAGCTCCCTTCCAG-3’ and reverse 5’-GGTACCCCGACACGGCAC-3’. The Luc2 sequence was amplified using the following primers: forward 5’-GGTGTGGAAAGTCCCCAGGC-3’ and reverse 5’-CACACAAAAAACCAACACAC-3’.

As a negative control we used the pCS2+ plasmid.

For the expression of human Alk5 dominant negative we used the previously established plasmid pCMV5B-TGFß receptor I K232R (Addgene) (Wieser, et al., 1995).

For the expression of zebrafish Alk5a, to define the baseline activation with the zebrafish protein in the human cells and TGFß ligand, we generated the pCMV:alk5a plasmid by inserting the *alk5a* coding sequence (NM_001037683.2) downstream of the CMV promoter in the pCMV-Tol2 plasmid (Addgene). The full-length *alk5a* coding sequence was amplified using the following primers: forward 5’-ATGAGCTCCGCCGCGATGG-3’ and reverse 5’-TTATATCTTGATGCCCTCTT-3’. For the validation of the zebrafish dominant negative Alk5a we generated the pCMV:DNalk5a-linker-GFP plasmid by cloning the truncated sequence of *alk5a* coding sequence downstream of the CMV promoter (Addgene), followed by a flexible linker and the GFP coding sequence. The *DNalk5a* coding sequence was amplified with the primers mentioned in the previous section. The Flexible linker + 5’GFP sequence was amplified using the following primers: forward 5’-CTTGGACCTGGACTCGGATCCGGAGTGAGCAAGGGCGAGGAGCT-3’ and reverse 5’-TTACTTGTACAGCTCGTCCA-3’.

HEK-293T cells, plated in a 24-well plate, were transfected with 300 ng/well p3TP:Luc2, SV40:hRLuc2 and 300 ng/well of another construct according to the well and 1.5 μl/well Lipofectamine 2000 Transfection Reagent (Thermo Fisher Scientific), for 5-6 hours, in 1xDMEM+Glutamax (Thermo Fisher Scientific) containing 10% FBS Superior (Biochrom) without antibiotic. Plasmids used are described above. The medium was then changed to 1xDMEM+Glutamax with 10% FBS and 1% Penicillin-Streptomycin (PenStrep, Sigma) and the cells were cultured over-night.

The next morning, the media were replaced with fresh serum-free media (with 1% PenStrep) containing 2 ng/ul TGFβ1 ligand (from a 10μg/ml stock), or no addition. After 24 hours, cells were rinsed in PBS and lysed with PLB Buffer for Luciferase Assay (Promega). The cell lysates’ supernatant was used to perform the luciferase assay, using the Dual-Luciferase^®^ Reporter Assay System (Promega), following manufacturer’s instructions.

Each experiment was carried out in duplicate (two wells per condition) in at least two independent experiments.

#### Ablation, drug treatments and heat-shock

Cellular ablation was induced with a 6-hour treatment in Mtz solution. Briefly, Mtz (Sigma) was eluted in DMSO to a concentration of 2.5 M immediately before treatment, and then diluted to a final concentration of 2.5 mM in fish system water. Fish were treated in individual containers and were kept in the dark during treatment to prevent degradation of Mtz. After exposure to Mtz, water was changed twice to remove any traces of drug and fish were kept in an incubator at 28°C until sacrifice or 7 dpab, when they were transferred to the system.

Treatment with the TGFß inhibitor SB431542 (Calbiochem) was adapted from previous reports (Jazwinska, et al., 2007; Chablais and Jazwinska, 2012). The compound was eluted in DMSO to a concentration of 15 mM and diluted in fish water to a final concentration of 15 μM. Fish were treated with the drug solution during the 24 hours prior to Mtz treatment and from 24 to 48 hpab.

To perform EdU incorporation, each animal was injected intraperitoneally with 20 μl of 10 mM EdU (Thermo Fisher) solution 3 hours prior organ collection. The stock solution was prepared in DMSO to a concentration of 1 M and diluted in PBS prior injection. Injections were performed using 29G U-100 insulin syringes (BD Micro-Fine).

Cre^ERT2^-based cell recombination was induced with intraperitoneal injection of 15 μl of 1.25 mM 4-hydroxytamoxifen (Sigma) diluted in PBS from a stock solution of 25 mM eluted in ethanol (Kikuchi, et al., 2010). Prior dilution, stock solution was heated at 65°C for 5 minutes, to ensure the complete dissolution of the reagent. Animals were injected for three consecutive days and let to recover in isolated recipients for two days prior cell ablation.

For heat-shock treatments, fish were placed in pre-heated water at 33°C for a period of 1 hour at 20 hpab and transferred to an individual container with system water at 28°C.

#### Cell transplants

For efficient engraftment of GFP+ WKM donor cells the host hematopoietic tissue must be ablated (Traver, et al., 2004). Therefore, *Tg(nfatc1:NTR-mCherry)* hosts were irradiated prior WKM transplantation. Fish were placed in petri dishes and irradiated with a single 15 Gy dose in a RS-2000 X-Ray irradiator (Rad Source) the two days before WKM transplant. WKMs were dissected from *Tg(ubi:switch);Tg(nfatc1:NTR-mCherry)* fish, placed in cold PBS + FBS 5%, and dissociated with a pipette. Cells were filtered through a 70 μm cell strainer (Corning) and centrifuged for 8 minutes at 300 x g. They were resuspended in 30 μl of 1x PBS per kidney and any trace of FBS was removed. A volume of 10 μl of cell suspension was injected intraperitoneally in each host. Injected animals were kept in the incubator for 7 days in clean water in individual containers to allow recovery and then transferred to the system. One month post transplantation, after reconstitution of the host kidney marrow, incorporation of the donor cells was confirmed by detection of GFP+ cells and the fish were subjected to valve cell ablation protocol as described above.

#### Doppler Echocardiography

Zebrafish were anesthetized using 0.016% buffered Tricaine diluted in system water. Animals were placed in supine position in a bed made of modelling clay, adjustable to the size of the fish. Fish were fully submerged in anesthesia solution to ensure propagation of the ultrasound signal. Vevo2100^®^ Imaging System (VisualSonics) and VisualSonics Ultrasound Imaging Software (Version 1.6.0) were used for echocardiography together with the high frequency MicroScan transducer (MS700 v3.0) at a frequency of 40MHz. Transducer head was oriented along the longitudinal axis of the fish and image was acquired with anterior side to the left and posterior side to the right. The imaging was performed in Color Doppler mode. To account for variations, videos were recorded for 10 seconds and across different 2D planes spanning the AV canal. Image acquisition was completed within 5 minutes of sedation to avoid cardiac function aberrations. After imaging, the fish were returned to a container containing system water and observed until recovery. Vevo Lab™ software package v.1.7.0 (VisualSonics) was used for image analysis.

#### Transmitted Electron Microscopy

Zebrafish hearts were dissected and fixed in 4% paraformaldehyde with 2.5% glutaraldehyde in 0.05 M HEPES buffer (pH 7.2) for 2 hours at room temperature, and subsequently stored at 4°C. To facilitate the orientation of the samples, hearts were pre-embedded in 3% LMP-Agarose. Samples were rinsed three times in 0.05 M HEPES buffer (pH 7.2) and post-fixed in 1% (w/v) OsO4 for 1 hour. After washing three times with distilled water, blocs were stained with 2% uranyl acetate for 1 hour. Samples were dehydrated through a graded series of ethanol washes, transferred to propylene oxide and embedded in Epon according to standard procedures (Laue, 2010). Tissue semi-thin sections (900 nm thick) were obtained in a Ultracut E microtome (Reichert-Jung, Leica) and stained with Richardson staining solution (Richardson, et al., 1960). Ultra-thin 70 nm sections were then collected on copper 2×1 slot grids. Sections were examined with a JEM-1400 Plus transmission electron microscope (Jeol, Japan), operated at an accelerating voltage of 120 kV. Digital images were recorded with an EM-14800Ruby Digital CCD camera unit.

#### Histology and Imaging

Hearts were fixed in 4% buffered paraformaldehyde for 1 hour at room-temperature, washed in 1x PBS and embedded as previously described (Mateus, et al., 2015). Briefly, the tissue was placed overnight at 4°C in a solution of 30% (w/v) sucrose prepared in 1x PBS, pre-embedded in 7.5% (w/v) porcine gelatin (Sigma)/15% (w/v) sucrose in 1x PBS at 37°C for 1 hour and embedded with a new solution of gelatin. Tissue blocks were frozen in isopentane (Sigma) cooled in liquid nitrogen. Cryosections were cut at 10 μm using a Leica CM3050S cryostat (Leica) and were kept at −20°C until further use. When preparing samples for *in situ* hybridization, all solutions and materials were kept RNase-free and slides were stored at −80°C.

Prior further analyses, slides were thawed for 10 minutes at room temperature and gelatin was removed in 1x PBS at 37°C. Immunodetection started with a wash in 0.1M glycine (Sigma) followed by permeabilization for 7 minutes at −20°C in pre-cooled acetone. Sections were incubated in a blocking solution of PBDX (1% (w/v) Bovine Serum Albumin, 1% (v/v) DMSO, 1% (v/v) Triton-X100 in PBS) with 15% (v/v) goat serum for a minimum of two hours at room temperature. Incubation with the following primary antibody was performed overnight at 4°C: GFP (1:200), mCherry (1:100), Elastin1 (1:100), Biotinylated-HABP (1:100), Fli1 (1:100), L-Plastin (1:200), pSmad3 (1:100). Slides were washed several times with PBDX and incubated with the corresponding Alexa Fluor conjugated secondary antibodies (1:500) overnight at 4°C. For all incubations, slides were covered with a piece of Parafilm-M to ensure homogenous distribution of the solution. Slides were washed a minimum of 3 times for 15 minutes each in a solution of 0.3% (v/v) Triton-X100 in PBS (PBST) and counterstained with 0.0002% (w/v) DAPI (Merck) in PBST for 10 minutes. Slides were then washed a minimum of 3 times for 15 minutes each in PBST and mounted with DAKO Fluorescence mounting medium.

Elastin1 antibody was purified from the previously described serum stock (Miao, et al., 2007). pSmad3 and Fli1 detection required an antigen retrieval step in 10mM Sodium citrate buffer pH 6.0 with 0.05% (v/v) Tween for 10 minutes at 95°C. HABP detection was adapted from a previous report (Govindan and Iovine, 2014) and included an additional step in methanol for 10 minutes at room temperature before the glycine.

For EdU detection, we followed the Click-iT EdU Cell Proliferation Kit for Imaging, Alexa Fluor 647 dye protocol (Thermo Fisher) and incubated each slide with 100 μl of Click-iT reaction solution for 30 minutes at room temperature prior incubation with antibodies.

For TUNEL detection, we followed the instructions from the In situ Cell Death Detection Kit, Fluorescein (Roche). We added a step in methanol for 10 minutes before the glycine and incubated each slide with 50 μl of TUNEL solution at 37°C for 1 hour, after incubation with secondary antibody.

*In situ* hybridization was performed following the RNAscope Fluorescent Assay (Advanced Cell Diagnosis) protocol for frozen tissue. We used a custom made *tgfβ1b* probe (Advanced Cell Diagnosis) and, when required, prior DAPI staining, we started the immunodetection protocol from the blocking solution step as described above.

All sections were imaged using a Zeiss LSM800 (Zeiss) inverted microscope, and ZEN 2.3 (Blue edition).

#### Data analyses and quantification

Image analyses and quantifications were performed using the FiJi ImageJ softawre (Schindelin, et al., 2012).

All quantifications represent an average of 3-5 serial mid-luminal sections per individual. Percentage of surviving cells was quantified by dividing the number of DAPI+ *nfatc1*+ cells inside the AV valve leaflets after ablation with an average (n=4) of the total number of *nfatc1*+ cells in uninjured AV valves. TUNEL+ cells represent the percentage of apopototic cells over the total number of DAPI+ *nfatc1*+ in that specific time point. The ratio El1/HABP was calculated from the Elastin1 and HABP total areas. pSmad3+ cells were counted in the AV canal region, excluding epicardium, myocardium and non luminal endocardium.

All statistical analyses were performed in GraphPad Prism (Version 6.07) and illustrations were done in Inkscape (XQuartz X11).

#### Laser Capture Microdissection

Hearts from uninjured, 48 hpab and 21 dpab animals were directly embedded in Tissue-Tek O.C.T. compound (VWR) without tissue fixation and were kept at −80°C until further use.

Cryosections with a thickness of 10 μm were placed on membrane coated glass slides (Leica) and were prevented from thawing until microdissection. Sections were rapidly dehydrated and briefly stained with Mayer’s hematoxylin and Bluing reagent (Scott’s tap water) for 15sec followed by washes in increasing gradient of 70 to 100% ethanol. AV valves and surrounding tissues were microdissected under optical control using the Laser microdissection device LMD6000 (Leica). The microdissected material was captured in Eppendorf tubes filled with 50 μl RNA lysis buffer (RLT+β-mercaptoethanol). We collected 3 biological replicates per time point, comprising a pool of 50-105 sections from 2 hearts per each replicate. After sample collection, the dissected material was preserved at −80°C until isolation of the RNA.

#### Transcriptome analysis

For RNA-seq, RNA was isolated from the laser microdissected samples using the miRNeasy micro Kit (Qiagen) combined with on-column DNase digestion (DNase-Free DNase Set, Qiagen) to avoid contamination by genomic DNA. RNA and library preparation integrity were verified with a BioAnalyzer 2100 (Agilent) or LabChip Gx Touch 24 (Perkin Elmer). 20 ng of total RNA was used as input for PolyA enrichment (NEXTflexTM PolyA Beads) followed by library preparation using NEXTflexTM Rapid Directional qRNA-SeqTM Kit (Bioo Scientific Corp). Sequencing was performed on the NextSeq500 instrument (Illumina) using v2 chemistry, resulting in average of 38M reads per library with 1×75bp single end setup. Raw reads were assessed for quality, adapter content and duplication rates with FastQC (Andrews S. 2010, FastQC: a quality control tool for high throughput sequence data. Available online at: http://www.bioinformatics.babraham.ac.uk/projects/fastqc). Trimmomatic version 0.33 was employed to trim reads after a quality drop below a mean of Q15 in a window of 5 nucleotides (Davis, et al., 2013). Only reads of at least 15 nucleotides were cleared for subsequent analyses. Trimmed and filtered reads were aligned versus the Ensembl zebrafish genome version danRer11 (GRCz11) using STAR 2.6.0c with the parameters “--outFilterMismatchNoverLmax 0.1 --alignIntronMax 200000” (Dobin, et al., 2013). The number of reads aligning to genes was counted with featureCounts 1.6.0 from the Subread package (Liao, et al., 2014). Only reads mapping at least partially inside exons were admitted and aggregated per gene. Reads overlapping multiple genes or aligning to multiple regions were excluded. Differentially expressed genes were identified using DESeq2 version 1.18.1 (Love, et al., 2014). Genes were classified to be significantly differentially expressed (DEG) with Benjamini-Hochberg corrected P-Value < 0.05 and - 0.59≤ Log2FC ≥+0.59. The Ensemble annotation was enriched with UniProt data (release 12.04.2018) based on Ensembl gene identifiers (Activities at the Universal Protein Resource (UniProt)).

MA plots were produced to show DEG regulation per contrast based on DESeq2 normalized counts. Dimension reduction analyses (PCA) were performed on DESeq2 normalized and regularized log transformed counts using the R packages FactoMineR and factoextra. For the assembly of the heatmap, DESeq normalized counts of all DEGs were transformed to a Z-score per row and clustered using k-means clustering (four clusters). Genes of each cluster were subjected to gene set enrichment analysis using KOBAS (Xie, et al., 2011). Main overrepresented pathways per cluster were identified and agglomerated based on manual inspection of significant Reactome database gene sets with P-Value <= 0.05.

#### Quantitative PCR

For qPCR analyses single hearts were homogenized with the Bullet Blender Gold (Next Advance). Total RNA was isolated using the RNeasy Mini Kit (Qiagen) followed by DNAse digestion (Invitrogen). cDNA was synthesized using the High Capacity RNA-to-cDNA kit (Applied Byosystems) and qPCR was performed in a BIO-RAD CFX Connect Real-Time System with the DyNAmo Color Flash SYBR Green Master Mix (Thermo Scientific).

